# Quantitative and unbiased lung alveolar septum assessment in a LPS experimental mouse model using 2D-spatial correlation image analysis from hematoxylin and eosin slides

**DOI:** 10.1101/2025.07.24.666548

**Authors:** Micaela Lopassio, María José García, Leonel Malacrida

## Abstract

Quantitative assessment of lung tissue architecture is essential for evaluating disease progression in experimental models of acute lung injury (ALI). However, conventional methods for measuring alveolar septum (tabique) thickness rely on manual annotation, which is time-consuming and observer-dependent, compromising comparability and reproducibility. In this study, we introduce a novel, fully automated approach based on two-dimensional spatial autocorrelation function (2D-ACF) analysis to quantify septum thickness from hematoxylin and eosin (H&E)-stained lung tissue sections.

After validating the method with a simulated data set, the new approach was applied to a murine model of Acute Lung Injury (ALI) induced by intratracheal instillation of lipopolysaccharide (LPS), under two dietary conditions: normal (ND) and high-fat diet (HFD), thereby testing our approach with a double hit protocol for ALI development. The 2D-ACF analysis provided a robust metric of structural organization, allowing for an unbiased estimation of mean septum thickness across entire tissue images. Compared to controls, LPS instillation increased septal thickness more than six-fold, with further thickening observed in the HFD+Instilled group. These findings were consistent across >400 images and captured subtle additive effects of metabolic stress on lung injury.

Traditional manual measurements of septum thickness exhibited substantial inter-observer variability, particularly in the LPS-instilled group, where structural heterogeneity made consistent interpretation challenging. This subjectivity limits the reproducibility of histopathological evaluations, especially under pathological conditions with subtle anatomopathological differences. In contrast, the 2D-ACF method offered a standardized and observer-independent approach. A direct comparison between methods revealed a strong and statistically significant correlation in the instilled group (r = 0.89, p < 0.001), indicating that the 2D-ACF captures key structural features aligned with expert assessments while reducing user bias.

In summary, the 2D-ACF framework offers a powerful alternative to conventional image analysis for studying lung pathology. Its adaptability to different experimental settings and potential to be extended to other tissues with spatial heterogeneity makes it a valuable tool for translational biomedical research and digital pathology.

## Introduction

Acute Respiratory Distress Syndrome (ARDS) and Acute Lung Injury (ALI) are major contributors to severe respiratory failure, with mortality rates between 30% and 60%, as reported in various studies [1,2]. These conditions frequently occur in Intensive Care Units (ICUs) and are among the primary reasons for requiring mechanical respiratory support in critical care patients.

The use of animal models has been crucial for understanding the mechanisms underlying lung injury. Among the various experimental models, the mouse model is widely used due to the availability of specific reagents and the ability to generate transgenic and “knock-out” mice, which help to understand molecular aspects of different pathologies [3]. The American Thoracic Society identifies key features of ALI in experimental models, such as tissue injury and disruption of the alveolar-capillary barrier. Assessing lung injury in ALI animal models requires careful and expert analysis of histological findings, which can be intensive and challenging to compare. A quantitative evaluation of various histological features is crucial for understanding the level of lung damage. Major indicators of lung injury include neutrophil accumulation in the alveolar or interstitial spaces, hyaline membrane formation, presence of proteinaceous debris like fibrin strands in the alveolar space, thickening of the alveolar wall (known as alveolar septum or tabique thickness), and alveolar collapse, characterized by a reduced air-to-tissue ratio [4–5]. Traditionally, these measurements are performed manually using stereological estimation techniques. This tool is based on morphometric methods that allow the extraction of three-dimensional quantitative data from two-dimensional elements. It uses intrinsic geometric properties of the studied structure, such as the alveolus in this case. These methods allow for determining cell numbers, the volume fraction occupied by an element, the length of an element, and the area of a structure by utilizing a grid integrated into the microscope eyepiece, which contains points, lines, planes, or volumes, depending on the cellular and tissue structures being studied [6,7]. For example, using the point-counting technique, it is possible to determine the air/tissue ratio in the alveolar region. By placing a grid with points on the microscope eyepiece, the number of points corresponding to tissue intersections and those corresponding to air are counted, and the ratio is established [8,9]. In this case, the measurements are reduced to counts, and while these techniques have multiple applications, they are not very precise (not to mention the time consumed to perform this analysis). The method is observer-dependent, and it is worth noting that it does not involve the digitalization of the analyzed image, which can lead to systematic bias affecting the estimation.

Optical microscopy enables the straightforward digitalization of hematoxylin-eosin histological sections, facilitating the quantification of areas and volumes in pixel units (e.g., μm²). Whole-slide scanners, for example, enable the relatively good resolution of a complete section [10,11]. Additionally, open-source software such as ImageJ or commercial tools like Image-Pro Plus offer various possibilities for semi-automatic digital data analysis [12]. In this case, at least 10 septum per image are estimated by using the drawing tool of FIJI, allowing for an average septum width per field. The same is done in 10 other fields per sample, generating a value for each histological piece under study. Despite the digitalization of images, analysis remains labor-intensive and prone to user bias.

Image correlation spectroscopy (ICS) is a quantitative image analysis technique based on the spatial autocorrelation function (ACF), which measures the degree of similarity between pixel intensities as a function of their spatial displacement [13,14]. By calculating how image intensity patterns repeat over space, ICS can reveal characteristic length scales and structural organization within a sample. Initially developed for fluorescence microscopy, ICS has proven particularly powerful in analyzing molecular mobility and aggregation in both fixed and live-cell imaging, owing to its ability to extract statistical descriptors from noisy, heterogeneous signals with minimal computational overhead [15, 16, 17, 18].

The ACF provides a reliable and unbiased method for analyzing spatial organization by integrating data across the entire image. This allows it to identify patterns and heterogeneities that might not be visible through visual inspection or threshold-based segmentation. ICS has been successfully applied beyond biological imaging, such as in geoscience, to assess the distribution of mineral phases in rock samples, demonstrating its broad applicability in measuring spatial heterogeneity in complex systems [19, 20, 21, 22].

This work proposes using the analytical power of ICS to evaluate pulmonary histopathology through the objective measurement of alveolar septum thickness in hematoxylin and eosin (H&E) stained lung sections. In diseases such as acute lung injury (ALI), the lung parenchyma undergoes remodeling that manifests as spatially heterogeneous changes in tabique thickness. ICS provides a scalable, unbiased, and statistically rigorous method to quantify these spatial features across entire tissue sections. The use of ICS represents a novel tool for unsupervised analysis of the spatial organization of lung tissue architecture in an experimental mouse model of ALI. By applying 2D spatial autocorrelation analysis to H&E images, we demonstrate ICS’s ability to capture and quantify variations in septum thickness in complex ALI models, thus offering a new avenue for characterizing tissue remodeling in lung pathology.

## Materials and Methods

### Experimental Model

C57BL/6 mice were used as the experimental model. The animals were included in the protocol starting at 8 weeks of age after verifying normal phenotypic traits. They were randomly divided into two groups, called the normal diet (ND) and the high-fat diet (HFD). The mice were fed ad libitum with their respective diets for 14 weeks and housed under specific pathogen-free conditions until two days before the experiment (at the animal facility of the Institut Pasteur de Montevideo). The mice were weighed weekly to monitor body mass gain and ensure roughly equal weight distribution between groups. A total of n=10 animals were used per group.

### LPS-Induced Lung Injury Model

An intratracheal instillation model of lipopolysaccharide (LPS) from Escherichia coli (O55 L2880, Sigma-Aldrich, St. Louis, MO) will be used, as previously employed by Files et al. [23]. C57BL/6 mice on either ND or HFD were anesthetized with isoflurane inhalation (100 mg/kg). After confirming an adequate anesthetic level, the animals were positioned at a 45° angle on a board for orotracheal intubation. A 20G catheter was inserted through the vocal cords, and 3 μg/g of LPS or an equivalent volume of saline (SF) was instilled. The mice received 10 μL/g of saline subcutaneously as preventive resuscitation, with free access to water and food.

This model is characterized by the development of sublethal acute lung injury (ALI), which peaks at 3-4 days and resolves progressively over a 10-day period. The animals were evaluated in terms of ventilatory mechanics (VM) at 4 days post-LPS [24]. The animal work was evaluated and approved by the National Ethics Committee with the animal protocol for Animal Use (N° 070153-000571-19).

### Histology

After measuring ventilatory mechanics, the lungs were extracted and fixed in formalin for 24 hours. The tissue was dehydrated in five steps of 30 minutes each with increasing concentrations of alcohol (50°, 70°, 96°, and isopropyl alcohol), followed by a 30-minute step in chloroform. The tissue was then embedded in paraffin, and serial histological sections were prepared (approximately 20 sections per slide). Pulmonary tissue was stained with hematoxylin and eosin.

### Image Acquisition

Tissue sections of lung tissue stained with hematoxylin and eosin were imaged using an Olympus IX81 microscope, equipped with an Olympus UPlanFL N 40x 0.75NA objective. A Kiralux 5.0 MP Color CMOS Camera with 8-bit depth was used. Thorlabs ThorCam software (v3.x) handled camera control and data acquisition. Ten random images were obtained for each histological tissue sample corresponding to the lung of a mouse. A total of 100 images were analyzed for each study group: Control group (Ctrl, C57BL/6 ND), Instilled group (Inst, C57BL/6 ND + LPS), High-Fat Diet group (HFD, C57BL/6 HFD), and HFD plus LPS instilled group (HFD-Inst, C57BL/6 HFD + LPS).

### Measurement of Alveolar Wall Thickness (Pulmonary Edema) Using the Traditional Method

Traditional analysis utilizes open-source software ImageJ (version 1.48v, Wayne Rasband, NIH, USA). The RGB images were first acquired and converted to 8-bit grayscale images. For measuring septum thickness (in cases of pulmonary edema), once the image was in this format, the “straight” tool in the toolbar was selected, which allows drawing a straight line across the width of the septum. The “Analyze-Measure” option was chosen to obtain the septum thickness value. This procedure was carried out for at least 10 septum in the same image, resulting in a table that expresses the values of all analyzed septum, which allowed us to obtain an average septum width per field. This value was then averaged across ten other fields per sample (Figure 1a and b).

**Figure 1:**
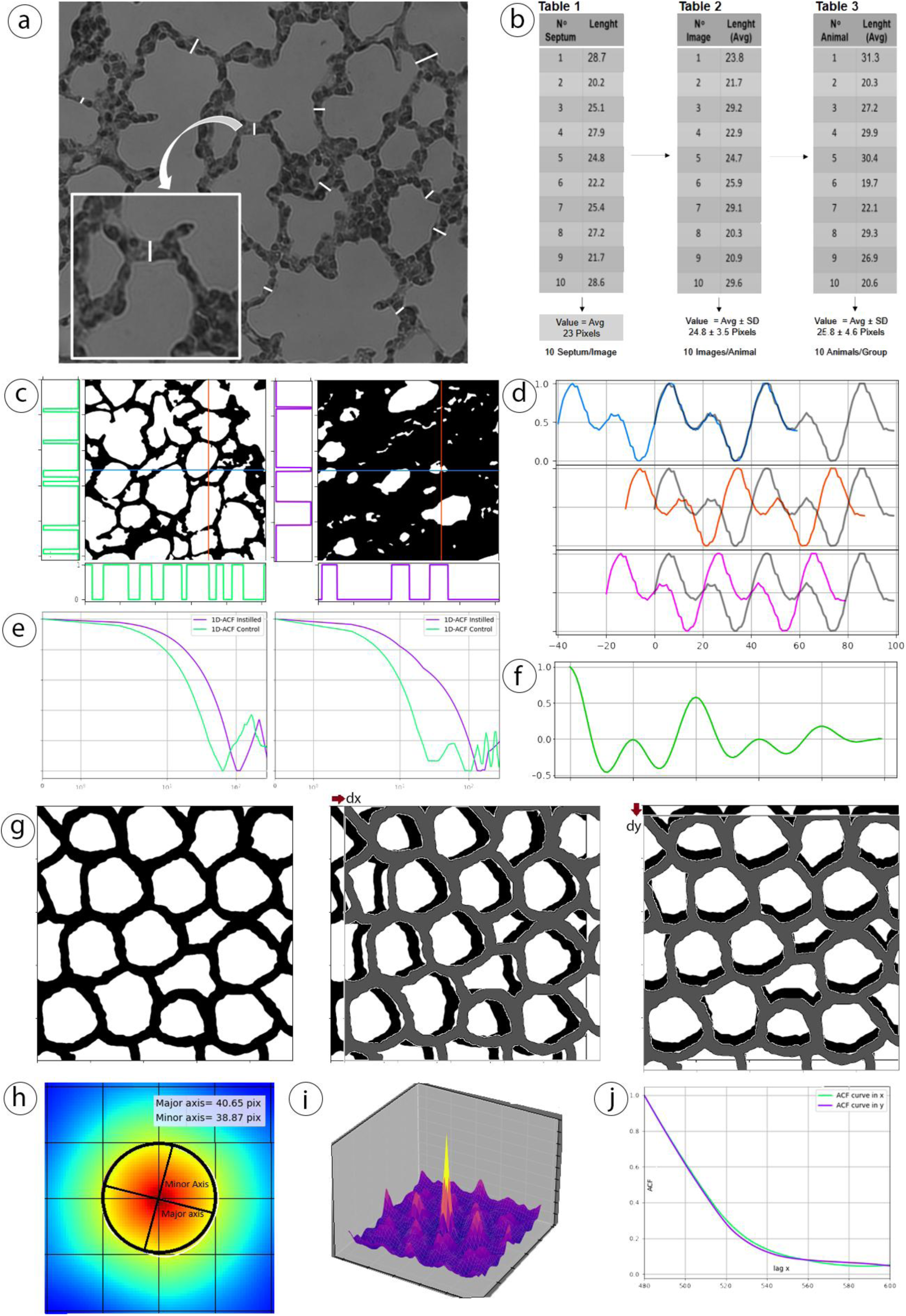
Autocorrelation-Based Quantification of Lung Septum Thickness. a) and b) Traditional method to quantify lung septum thickness. c) Mathematical explanation of autocorrelation function and its demonstration in the 1-dimensional case. A row and column selected for control and pathologic study groups after binarization, and its corresponding intensity plot level as a function of the pixel. e) 1-dimensional autocorrelation calculation for these rows and columns, in logarithmic scale, clearly shows the differences in the decay of the curve between both groups, noting that for the control group (green), the deviation is smaller compared with the pathological (violet). d) illustrative intensity signal and its shifted copies, in blue, orange and violet which translated distances T, T/4, T/2 respectively and the corresponding 1D ACF for that signal where it shows the correspondence between the peaks of ACF and the particular lags where the relationship between the curve and its translated version is maximal, opposite or cancels out, g) Simulated image to illustrate 2D sACF, the original image is moved x pixels to right and y pixels. h) Resulted 2D sACF from the previous image and fitted ellipse with its major and minor axis indicating the average size, i) Resulted 2D sACF surface in 3D, j) Central row and column plot of the 2D sACF.

### Python Code for the Calculation of Two-Dimensional Spatial Autocorrelation Function (2D-sACF)

A new method is proposed to identify pulmonary edema based on the 2-Dimensional Spatial Autocorrelation Function (2D-sACF). The technique was implemented using Python 3.12 as the programming language and utilizing code libraries such as OpenCV, scikit-learn, skimage, NumPy, Matplotlib, and SciPy, among others. The approach was applied to hematoxylin-eosin RGB images from lung tissue. To improve the algorithm’s efficiency the autocorrelation was calculated using the Wiener-Khinchin theorem, which states that the autocorrelation is the inverse Fourier transform of the power spectral density (PSD), which is the product of the Fourier transform of the image 𝐹{𝐼(𝑥, 𝑦)} and its complex conjugate 𝐹{𝐼(𝑥, 𝑦)}* [1]. Thus, the implemented method first calculates the 2D Fourier transform of the normalized image (image minus its average intensity value 𝐼̂), through the Fast Fourier Transform (FFT), then antitransforms the product of the transform by its conjugate and finally divides by the deviation 𝜎_𝐼_ of the image and its total size nxm.

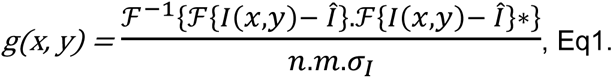

Since, after preprocessing, we got a binary image where pixel values are restricted to 0 and 1, normalization by subtracting the mean in this case is unnecessary. Typically, this correction is needed to standardize the intensity scale for images with a broader range of values, such as grayscale images. Here, the pixel values are already confined to a narrow range, and normalization would not improve the calculation of the ACF. In essence, the autocorrelation function describes how a signal matches with shifted versions of itself. As the shift increases, the function compares the signal with more distant regions, revealing how segments of the signal relate to one another.

To understand the fundamentals of autocorrelation, Figure 1 illustrates the concept in one dimension. In panel 1c, two representative images from the control and pathological groups are presented. In each image, a single row and a single column are arbitrarily selected, generating one-dimensional arrays of size 1 × n. For each of these one-dimensional signals, the autocorrelation function (ACF) is computed by shifting the signal to the right by a number of pixels d (referred to as the lag), multiplying each pair of corresponding pixel intensities from the original and the shifted signal, summing the products, and repeating this process for all possible shifts. When the lag is small, the values are highly similar, reaching a maximum of 1 at lag = 0, where the shifted signal perfectly overlaps the original. As the lag increases, the similarity decreases unless repeated patterns exist within the signal. The intensity profiles of these rows are shown, where it is evident that the control image exhibits shorter periods of low intensity (black = tissue), corresponding to a thinner alveolar septum, and longer periods of high intensity (white = air spaces), relative to the pathological (instilled) group. In the control sample, the frequency of signal transitions—decreases followed by increases—is higher, reflecting the higher prevalence of a narrow septum. On the other hand, the instilled sample displays fewer transitions and more prolonged low-intensity regions, indicating a thicker septum and increased tissue content.

Figure 1.e illustrates the resulting 1D ACFs from these profiles. In the control sample, the ACF decays more rapidly, reflecting the shorter correlation length associated with a thinner septum. In contrast, the slower decay observed in the instilled sample corresponds to a broader spatial distribution of tissue, i.e., a thicker septum, which is captured by the increased width of the ACF central peak.

To further clarify this concept, Figure 1.d shows a synthetic periodic signal with a known period T = 20 pixels. Colored versions of this signal are presented with shifts of T/4, T/2, and T pixels. The autocorrelation function g(d) is calculated by multiplying the original signal I(x) by its shifted version I(x−d) for each lag d, and summing the product across the entire signal length n. The resulting ACF shows distinct peaks at lag = 0 (maximum correlation) and lag = T (periodicity), while showing zero or negative values at T/4 and T/2, indicating no correlation or anticorrelation, respectively. These features correspond to the degree of overlap between the original and shifted signals. In this way, the ACF quantifies the probability that two pixels separated by a distance d have similar intensity values. By systematically comparing all possible pixel pairs at varying separations, the ACF provides a statistical measure of spatial similarity as a function of distance and direction [25].

Finally, subfigure 1.g extends this analysis to two dimensions. The same principles apply, now considering the image as a 2D matrix where each pixel holds an intensity value. The 2D ACF is computed by shifting the image in all directions, multiplying the original and shifted pixel intensities, and summing across the image. When structures within the image are small, the central peak of the ACF is narrow and decays rapidly with increasing shift. In contrast, larger structures result in broader central peaks and slower decay of the ACF, as is observed in the presence of pulmonary edema or septum thickening. Thus, the 2D ACF enables quantitative assessment of spatial organization in histological images and reveals tissue-level heterogeneity in a statistically robust manner.

The methods yield valuable information about the spatial characteristics of the image, including the distribution of different structures along its length and width, their shape and preferential orientation, and the various patterns that can be present [19]. The average size of the objects contained in it is of particular interest in this manuscript:

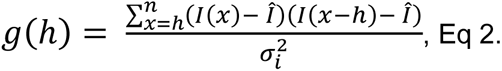

Equation 2 represents the formula for the calculation of the 1-dimensional Autocorrelation Function.

Before compute the 2D-sACF, each image is resized to size 1024x1024 pixels, and the following steps are applied, with the final goal of achieving a clean binary image, without background and without white gaps in the structure of the tissue, so that every region of the image that the human eye assigns as tissue, for the algorithm represents the same structure:

1. Convert image to grayscale.
2. Gaussian filter with kernel size 5x5 and sigma x=sigma y = 12.
3. Gaussian Mixture Model (GMM) segmentation technique with k = 3 (# of clusters).
4. Thresholding assigned a value of 0 to all pixels below the clusters’ highest intensity value.
5. Remove objects with a minimum size of 30 pixels to fill white holes in the structure.
6. Finally, fill holes to remove point marks in the white background corresponding to air.

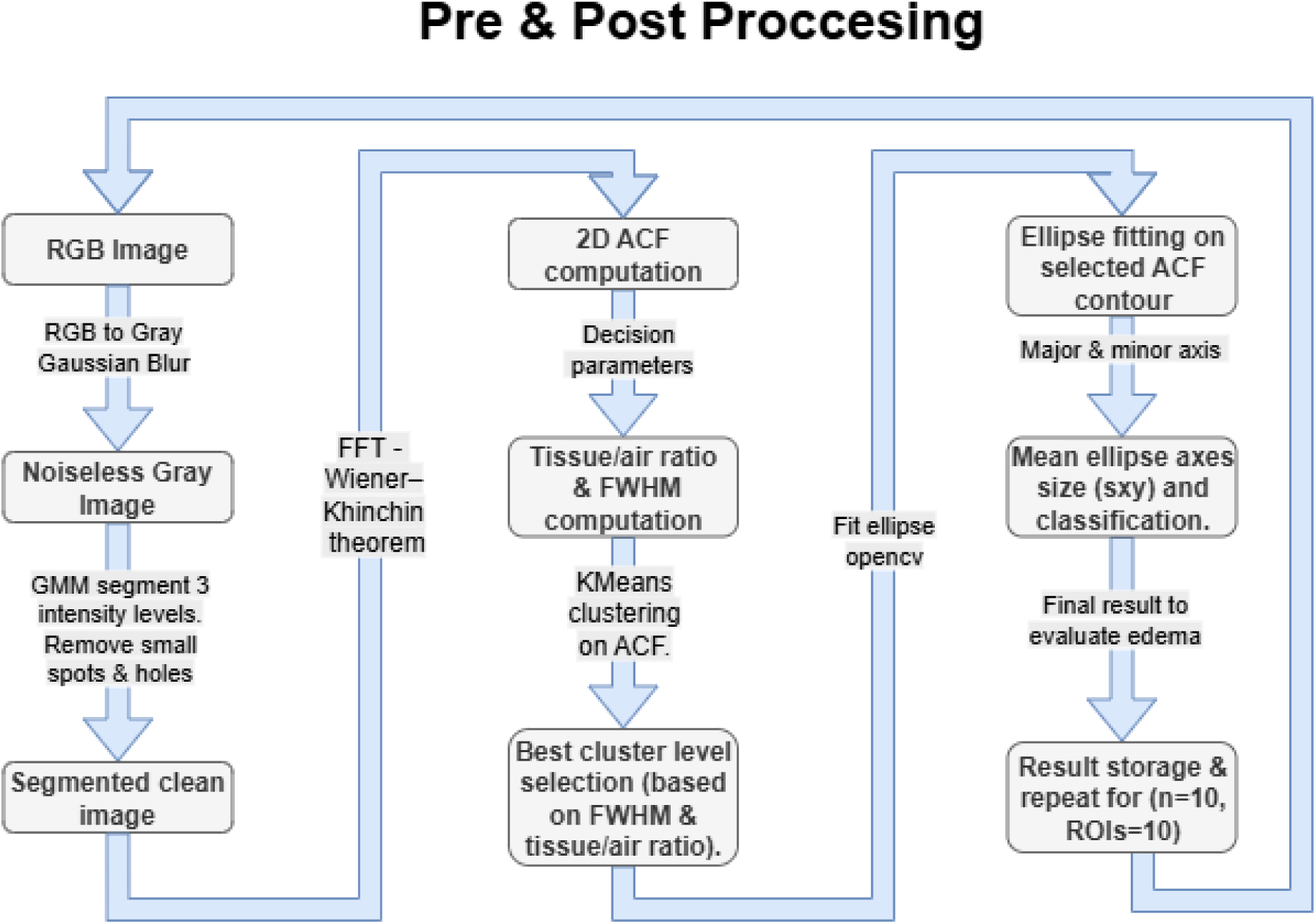

Each image processing parameter—such as the number of clusters used for segmentation, the size of the Gaussian filter, and the thresholding value—was chosen to strike a balance between preserving the tissue structure and avoiding artifacts that could artificially alter the measured area (e.g., artificially thickening or thinning the septum). The result was a binary image with only two intensity values, minimal background noise, and virtually no gaps in the segmented tissue structure.

To validate the effectiveness of this image preprocessing approach, four test cases were simulated using an image editor, based on a representative real image. These cases represent specific physiological or pathological conditions: (1) Normal alveolus without edema, (2) Overdistended alveolus without edema (super-alveolus), (3) Alveolus with edema, and (4) Overdistended alveolus with edema. These are shown in Figure 2.

**Figure 2.**
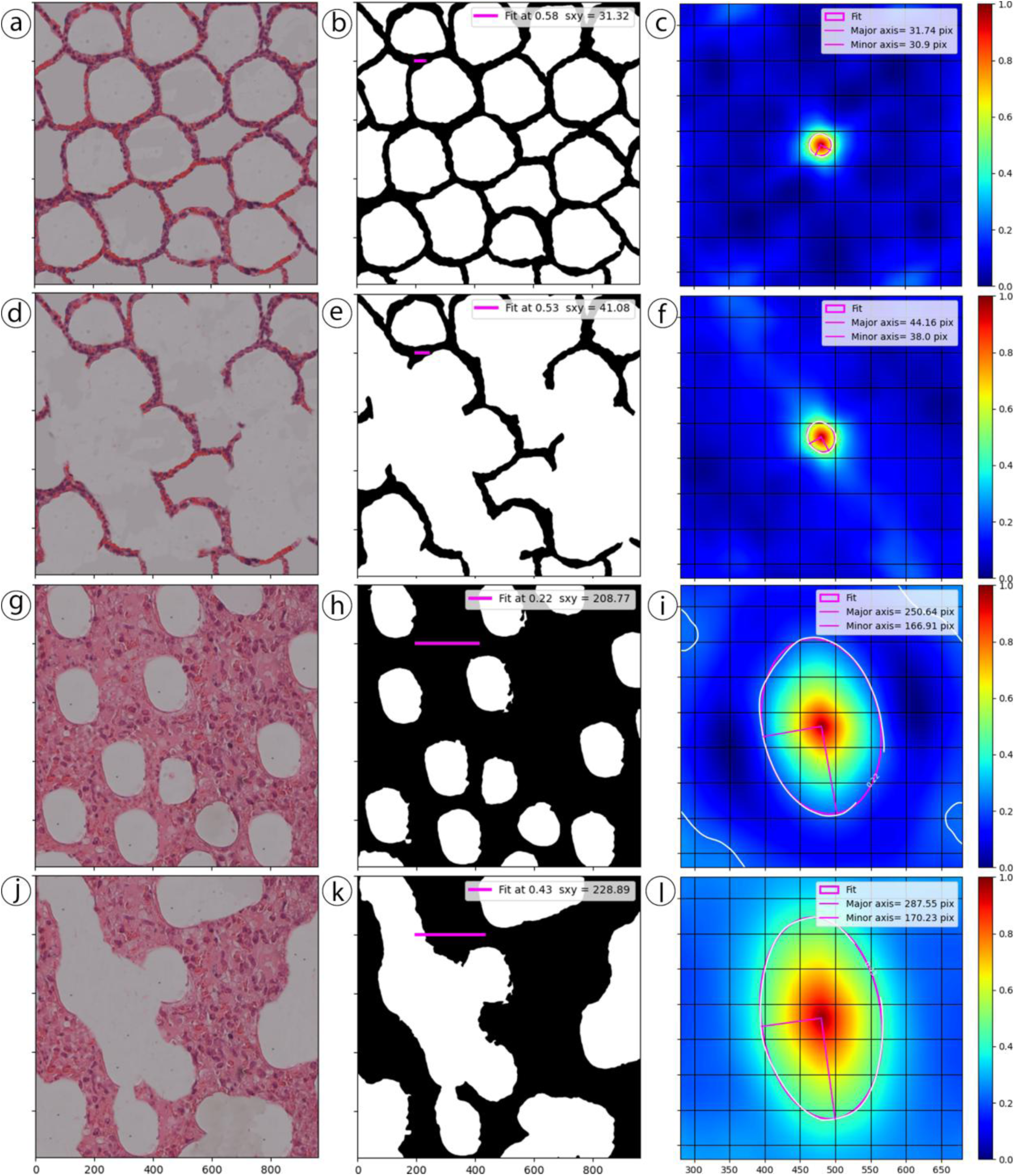
Simulated replica images were used to test the efficacy and robustness of the proposed 2D autocorrelation function (ACF) method in four defined conditions. Each image represents a simulation designed to mimic key structural characteristics observed in the four experimental groups: Control (a-c), Instilled (d-f), HFD (g-i), and HFD Instilled (j-l). These simulated images serve as ground truth for assessing the overall method of image segmentation and 2D-ACF analysis, utilizing ellipse fitting to measure characteristic tissue sizes.

To quantify the average size of the spatial structures in each image, a fitting procedure was applied to the two-dimensional spatial autocorrelation function (2D-sACF), which is assumed to follow an ellipsoidal bivariate Gaussian distribution. To identify the region of interest within the 2D-sACF, a dynamic threshold was computed using the K-means clustering algorithm, which partitions the autocorrelation data into K clusters based on similarity in intensity.

In early stages, the fitting was performed using Python’s lmfit library, based on a parametric bivariate Gaussian model. This model describes an elliptical shape defined by coefficients a, b, and c, which were initially estimated using the method of moments [26]. However, this approach was not robust across all images, as it depended heavily on selecting an optimal cutoff point and on assuming a specific functional form for the fit—both of which proved problematic.

To overcome these limitations, a non-parametric geometric fitting method was implemented. Recognizing that the shape of the 2D-sACF is approximately elliptical, the method involves thresholding the 2D-sACF at a chosen contour level and fitting the resulting shape directly to a geometric ellipse. This approach avoids the need for an explicit model or function, and it can be applied consistently across images. Using the OpenCV library’s fitEllipse function, the contour is fitted, and the major and minor axes of the ellipse—corresponding to the standard deviations σₓ and σᵧ of the autocorrelation function—are extracted. These parameters quantitatively describe the spatial correlation length in both directions.

Steps performed in the image analysis pipeline:

1. Compute the 2D spatial autocorrelation (2D-sACF) from the binary image.
2. Use peak_local_max from the scikit-image library to identify local minima and select the appropriate threshold.
3. Threshold the ACF matrix at the chosen level to extract the main peak region.
4. Fit the contour to an ellipse using cv2.fitEllipse from the OpenCV library.
5. Extract the ellipse’s major and minor axes (σₓ and σᵧ) as measures of the spatial correlation in the image.
6. Store the mean values of these axes for each image for subsequent statistical analysis.

A morphological opening operation (erosion followed by dilation) with a kernel size of 11 pixels was also applied as a final step to remove residual small artifacts (”spots”) from the segmented images.

### Statistical analysis

All statistical analyses were performed using JASP software (version 0.19.3), JASP Team, University of Amsterdam. Before hypothesis testing, the data distribution was assessed using the Shapiro–Wilk test to evaluate the assumption of normality (see tables S1 and S2 in the supplementary material). Based on the test results, either parametric or non-parametric methods were applied accordingly.

For comparisons between two independent groups, data that followed a normal distribution were analyzed using an independent samples t-test. When normality was not met in at least one of the groups, the Mann–Whitney U test was used as a non-parametric alternative.

For comparisons involving more than two independent groups, a one-way ANOVA was used when all groups met the assumption of normality. In cases where the ANOVA indicated a significant difference, Tukey’s honestly significant difference (HSD) test was employed for post hoc pairwise comparisons. When the assumption of normality was violated, the Kruskal–Wallis H test was applied as a non-parametric alternative. Upon observing significant group differences with the Kruskal–Wallis test, Dunn’s post hoc test was conducted to assess pairwise differences.

A significance threshold of p < 0.05 (two-tailed) was used for all statistical tests. Descriptive statistics and tables were also generated within JASP to aid in data interpretation and visualization.

## Results

### Evaluation of the 2D-ACF analysis in simulated images

To evaluate the performance of the new method, the 2D-ACF was tested using a set of simulated images that represent pathognomonic features in healthy and diseased lungs (1: Normal alveolus without edema; 2: Overdistended alveolus without edema; 3: Alveolus with edema; and 4: Overdistended alveolus with edema). Figures 2b, e, h, and k illustrate the performance of our segmentation pipeline in generating masks to distinguish between tissue and air. The protocol achieves strong performance in filling holes and effectively identifying tissue boundaries. The 2D-ACF for each simulated tissue was calculated, and the fitting results are shown in the insets of Figures 2c, f, i, and l. Note that the 2D-ACF is not always round; in fact, only the “control” image shows similar short and long axes (31.7 and 30.9 pixels). However, in the other three examples, a long axis of the 2D-ACF is always present (Figures 2f, i, and l). This indicates an anisotropic distribution of tabique thickness, with the tabique being thicker in one direction of the image. Additionally, the long and short axes of the 2D-ACF can provide an angle that reflects the overall orientation of the tabique thickness in the image. Although we did not explore this aspect in this manuscript, it is another value that can be derived from the fitting analysis of the 2D-ACF. To determine the overall tabique thickness, the average of the major and minor axes of the 2D-ACF was calculated. In Figures 2b, e, h, and k, the average tabique thickness is represented as a pink line in the images, demonstrating the protocol’s effectiveness in measuring the mean thickness in each image.

### Validation of the 2D ACF Method Across Experimental Groups

To further challenge the performance of the two-dimensional autocorrelation function (2D-ACF) approach for quantifying the tabique thickness in experimental data, the analysis was applied to four lung groups: Control (Ctrl), Instilled (Inst), High-Fat Diet (HFD), and HFD-Instilled (HFD+Inst). Representative RGB images of H&E for each group are presented in Figure 3a, d, g, and j; the analysis was performed on 10 images per animal. The 2D-ACF for each respective image is presented in Figure 3c, f, i, and l, showing increased radios for HFD and HFD+Inst. The fitting results for each 2D-ACF show a major (σᵧ) and minor (σₓ) axes. Then, an average spatial correlation length (σₓᵧ) is calculated as the average of the major and minor axes of the 2D autocorrelation function (ACF), which is used as a quantitative proxy for the structural organization of each image. Noticeably, all groups show an anisotropic 2D-ACF, highlighting the nature of pulmonary architecture. This analysis may serve as a powerful tool for quantifying structural remodeling in pathological lung states. Instillation (Inst) has a strong impact on the σₓᵧ, increasing the overall tabique thickness six times compared to the control (Ctrl). Moreover, the combination of HFD with instillation (HFD+Inst) doubled the overall tabique thickness (276.8 vs 235.4, HFD+Inst and HFD, respectively).

**Figure 3:**
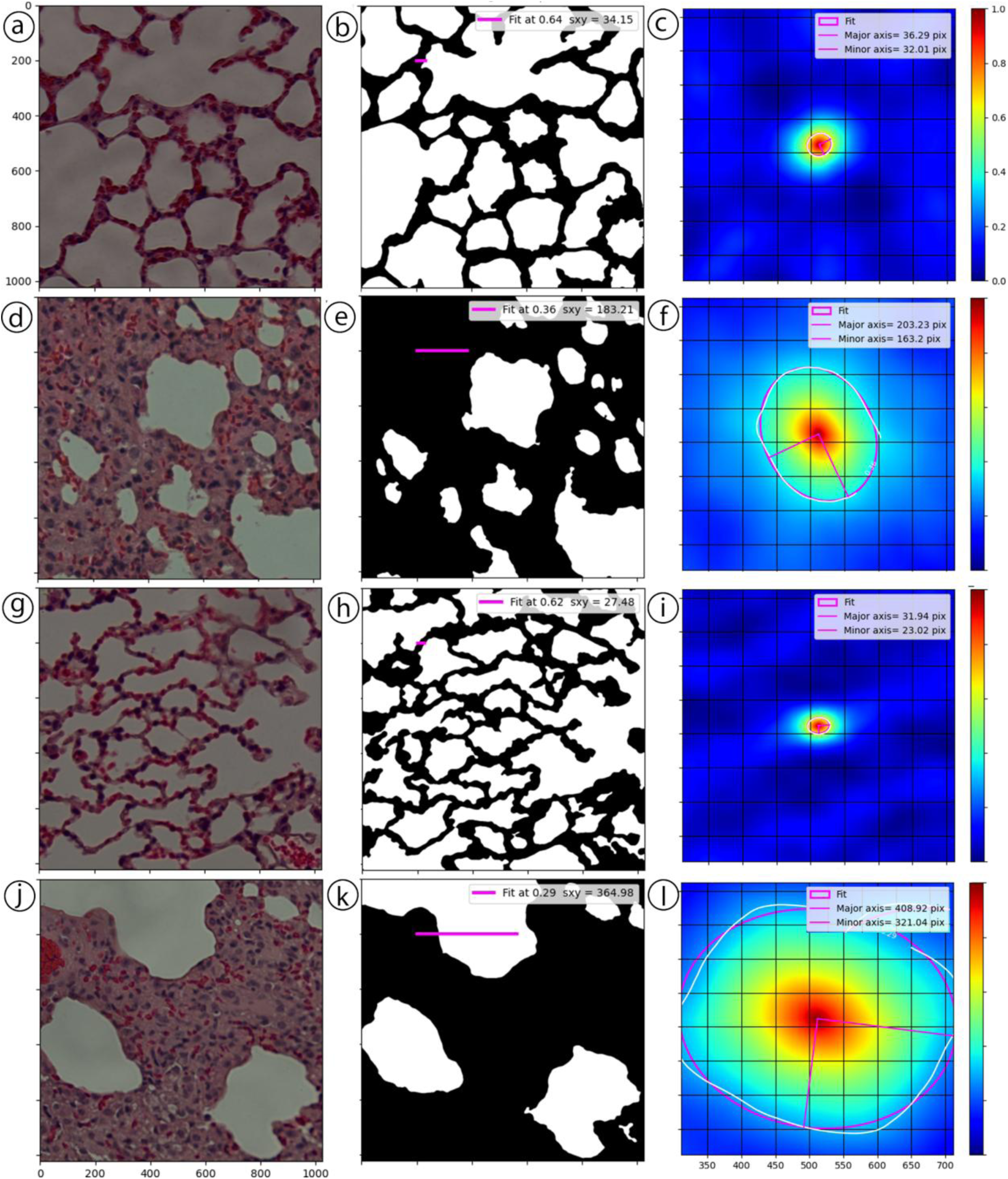
Demonstration and validation of the proposed 2d autocorrelation function method for quantifying edema and tissue structure across different experimental groups. Each row represents a different group: (a) Control, (d) Instilled, (g) HFD, and (j) HFD Instilled. Columns display: (a) the original histological image in RGB without preprocessing; (**b**) the processed binary image after applying multiple filtering steps to isolate tissue structures; and (**c**) the 2D spatial autocorrelation function heatmap in a jet colormap, with the fitted ellipse overlaid. The fitted ellipse quantifies the size of tissue structure, with its major and minor axes annotated. The control and HFD groups exhibit more uniform alveolar spaces, while the instilled and HFD-instilled groups show enlarged tissue and alveolar structures and higher heterogeneity.

The analysis of all the regions of interest (ROÍs) in each group yields the results presented in Figure 4. Notice that each segment in the violin plot represents 10 ROIs per animal; the results of all animals demonstrate two primary outcomes. First, the HFD has a basal background with alveolar tabique thickness increasing from 51.8±8.8 to 93.9±29.2 pixels (Ctrl vs HFD); this difference is statistically significant with a p-value < 0.05. The installation increased the overall tabique thickness by 235.4±24.8 pixels, and the combination of a high-fat diet and installation enlarged the overall tabique size even further to 276.8±48.7 pixels. Statistical analysis supports the group differences when Ctrl is compared to HFD and Inst, and when HFD+Inst is compared to Inst or HFD.

**Figure 4:**
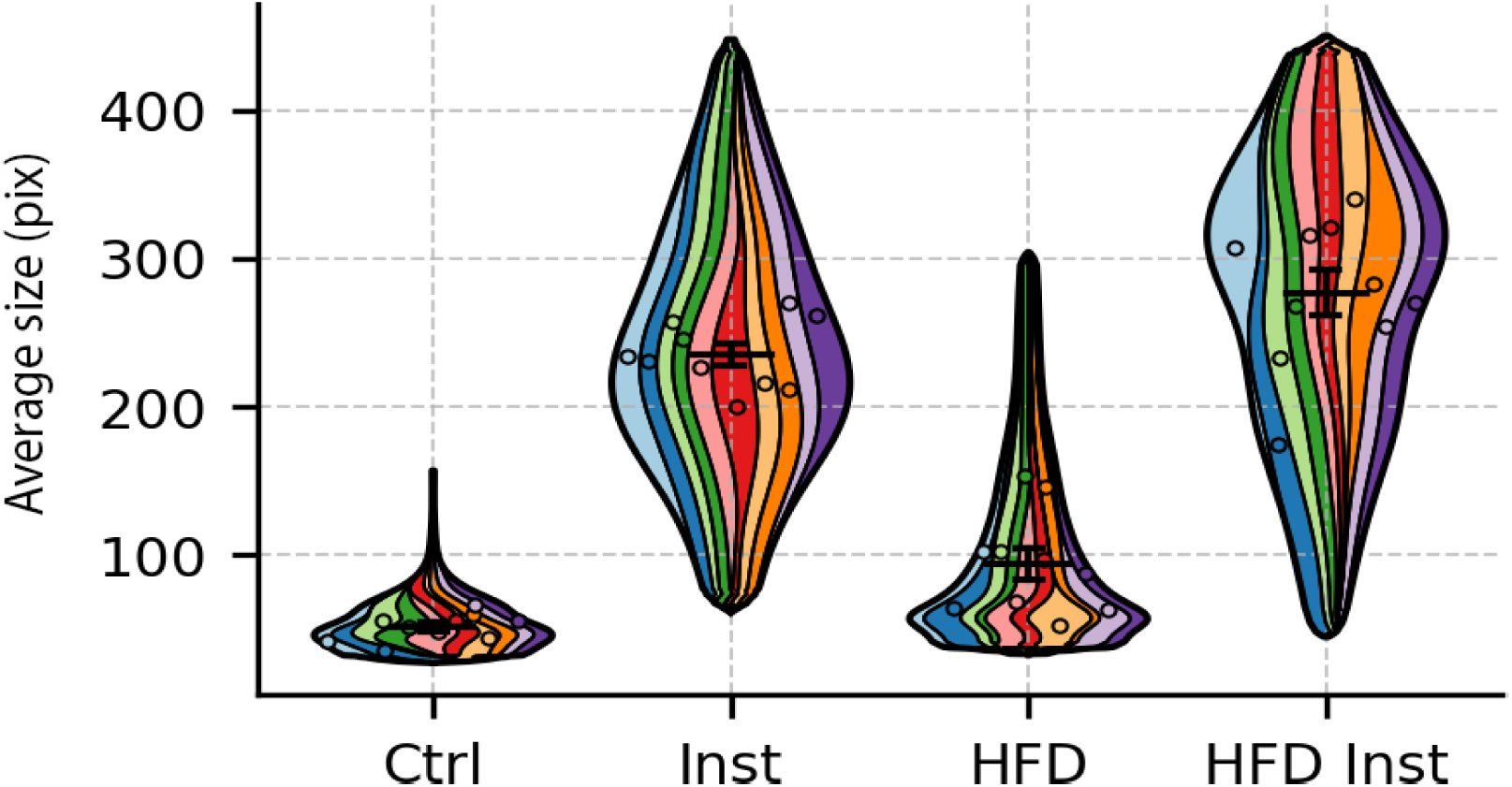
Super violin plots for the 2D-ACF result comparing the four groups of animals. The plot shows the average size distribution across the Control, Instilled, High-Fat Diet, and High-Fat Diet Instilled groups. Each plot combines a violin plot (showing the probability density), a box plot (showing the median and interquartile range), and individual data points (showing raw data distribution). The violin plots illustrate the probability density of the data at different values, providing insight into the variability and distribution shape within each group. The results reveal distinct differences between groups, with a broader spread in the Inst and HFD Inst groups, suggesting increased heterogeneity, and a higher distribution in the Instilled group, indicating a systematic change in the measured size.

### Tabique thickness quantification comparison between 2D-ACF and traditional measurement approach

Three independent analysts, blinded to experimental group assignments, quantified septal thickness using the traditional method on the same set of histological images. Each analyst performed the measurements independently. A summary of their results is presented in Figure 5 and Table 2. Notably, while all three observers consistently distinguished between the control and LPS-instilled groups, their reported mean septum thickness values showed considerable variability within each group. This discrepancy was particularly pronounced in the LPS-instilled group, where increased heterogeneity in alveolar structure and septal thickening likely contributed to divergent interpretations. Statistical analysis confirmed significant inter-observer variability, which poses challenges for reproducibility and highlights the subjective nature of manual histopathological assessment.

**Figure 5:**
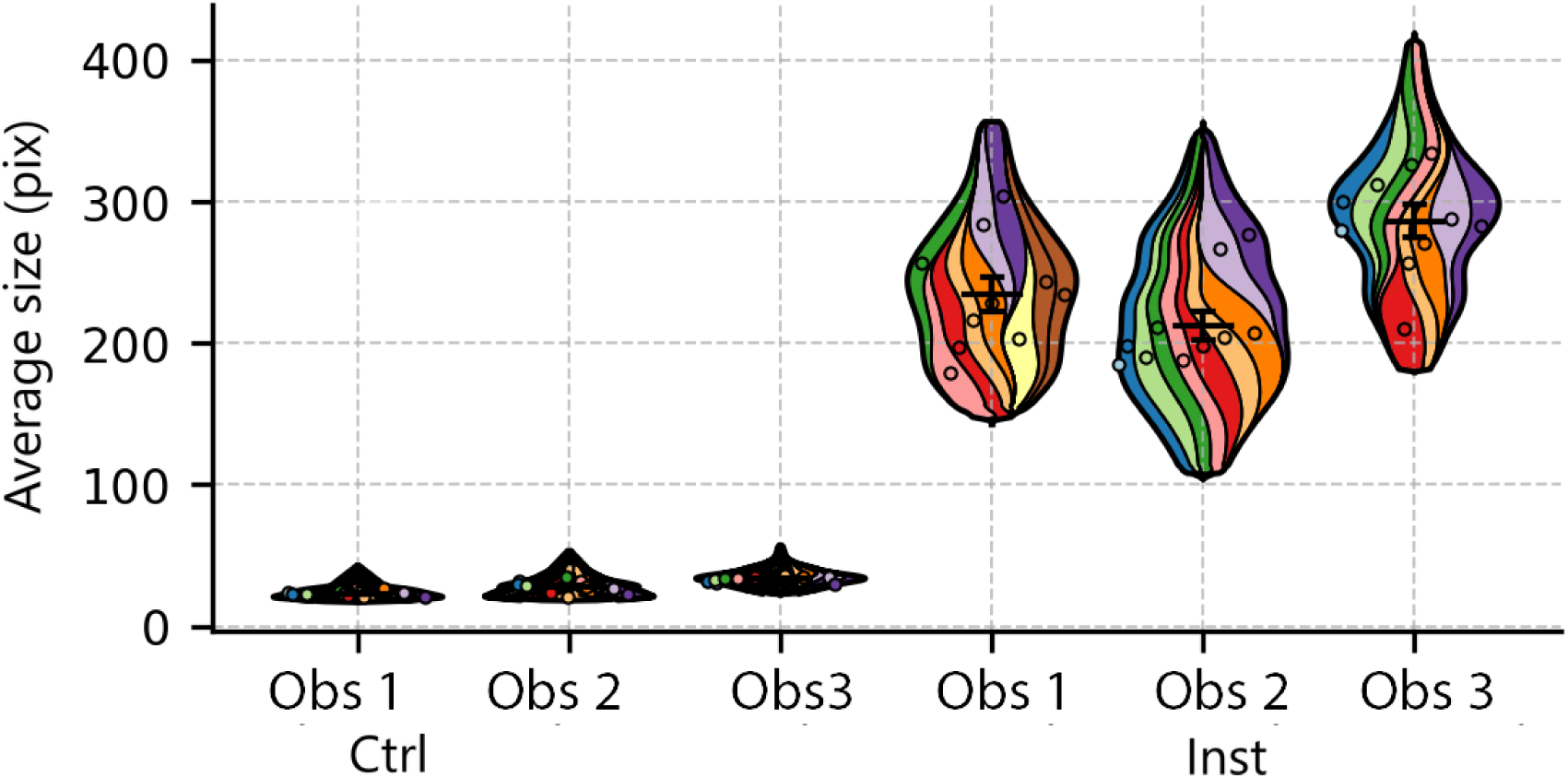
Violin plots for the results of three independent observers’ analysis of traditional tabique thickness. The measurements were obtained by three independent individuals (observers) using the conventional method. Each violin plot represents the density distribution of measurements recorded by each group, with embedded box plots showing the median, interquartile range, and individual data points. The results highlight the variability in manual measurements between observers, with observer 3 exhibiting a wider range of values, suggesting greater subjectivity, and observers 1 and 2 showing more consistency in their assessments.

To evaluate the agreement between the traditional and 2D-ACF methods, correlation analyses were conducted based on the average septal thickness measured per animal in both control and instilled groups (Figure 6a–b). In the instilled group, a strong correlation was observed between the two methods (r = 0.89, R² = 0.80, p < 0.001), indicating a high degree of consistency. In contrast, the control group exhibited a moderate correlation (r = 0.64, R² = 0.41, p = 0.046), reflecting increased variability in manual assessments of a thinner septum. These findings support the 2D-ACF approach as a reliable and reproducible alternative for quantifying septal thickness, particularly in conditions with substantial tissue remodeling.

**Figure 6:**
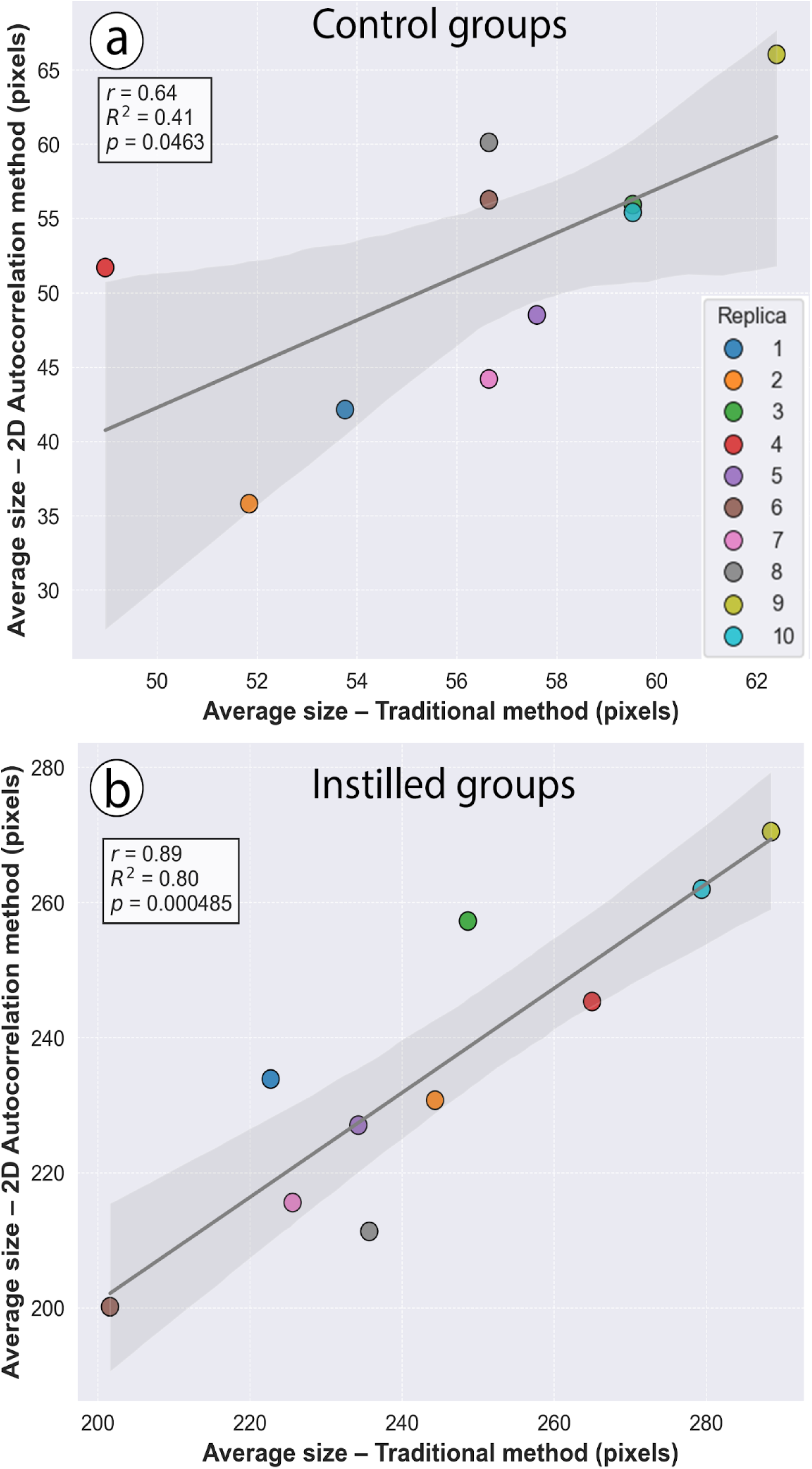
Correlation plots for traditional vs 2D-ACF analysis of tabique thickness. (a) Control group shows a moderate positive correlation between the two methods. (b) Instilled group exhibits a strong positive correlation. Each point represents the average measurement per animal (n = 10 per group). Gray shading indicates 95% confidence intervals for linear regression fits.

## Discussion

Histopathological evaluation of lung tissue remains a cornerstone for assessing disease progression in experimental models of acute lung injury (ALI). However, conventional analysis of lung slices is often qualitative or semi-quantitative, relying heavily on expert interpretation and thus subject to observer bias and variability [12, 24]. Moreover, manual assessments are time-consuming and lack scalability for high-throughput or longitudinal studies. In this context, the two-dimensional autocorrelation function (2D-ACF) approach presented in this study offers a novel, robust, and unbiased method for directly quantifying overall alveolar septum thickness from histological images. By leveraging spatial statistics [25], this technique overcomes the limitations of subjective evaluation, providing rapid, reproducible, and quantitative measurements of tissue architecture that advance the precision and efficiency of lung pathology analysis.

The animal model employed in this study was designed to emulate key pathological features of clinical ALI [3]. Intratracheal instillation of lipopolysaccharide (LPS) was used to induce lung injury [23], a well-established approach that reproduces many aspects of acute inflammation and alveolar damage [4, 28, 29]. Nonetheless, ALI in the clinical setting often occurs in patients with comorbidities such as obesity or metabolic syndrome, which are known to exacerbate lung injury [1,5]. To reflect this complexity, experimental groups were fed a high-fat diet (HFD), providing a more physiologically relevant background. The integration of these experimental conditions enabled us to examine how pre-existing metabolic alterations modulate the response to LPS-induced injury, revealing more pronounced tissue remodeling and alveolar septum thickening in the HFD-instilled group compared to the LPS-alone group (Figure 4 and Table 1).

**Table 1:** Descriptive comparisons of septum thickness measurements using the 2D-ACF method.

|  | Control | Instilled | HFD | HFD + Instilled |
| --- | --- | --- | --- | --- |
| **Mean ± SD** | 51.8 ± 8.8 | 235.4 ± 24.8 | 93.9 ± 29.2 | 276.8 ± 48.7 |
| Data are presented as mean ± standard deviation. All groups met the assumption of normality. Group comparisons were assessed using one-way ANOVA, followed by Tukey's post hoc test. Significant differences in septum (tabique) thickness between groups:<br>Control vs. Instilled: **** $p < 0.001$<br>Control vs. HFD: * $p < 0.05$<br>Control vs. HFD + Instilled: **** $p < 0.001$<br>Instilled vs. HFD: **** $p < 0.001$<br>Instilled vs. HFD + Instilled: * $p < 0.05$<br>HFD vs. HFD + Instilled: **** $p < 0.001$ | | | | |

One of the most significant limitations of traditional analysis is the dependence on manually selected septum regions. Figure 5 evidenced this situation, where inter-observer variability in measured septum thickness was statistically significant (see Table 2 and Figure 5). Observer 3 consistently produced significantly different measurements when compared to Observers 1 and 2 analyzing the Control or Instilled groups. This problem is particularly severe when comparing results between different observers within a group, and one can predict even more difficulties when comparing results between various research groups. The primary reason for this difference is the observers’ perception of what a septum is and how to assess the overall septum thickness based on the image under study (see Figure 3a, d, g, and j). As shown in descriptive Table 2, the variability in manual assessment is further highlighted by the percentage variation observed among the three observers. In the control group, the average inter-observer variation—calculated as the ratio of standard deviation to mean septum thickness—was approximately 9.97%, reflecting relatively consistent measurements in healthy lung tissue. In contrast, the instilled group exhibited a markedly higher variation of 14.75%, indicating greater subjectivity and difficulty in defining septum boundaries in lungs that are inflamed or structurally remodeled. These findings underscore the inherent limitations of manual analysis, particularly under pathological conditions where tissue architecture is disrupted.

**Table 2:** Descriptive comparisons of traditional septum thickness measurements by three independent observers.

| Group | Observer 1 | Observer 2 | Observer 3 |
| --- | --- | --- | --- |
| Control | 23.1 ± 1.8 | 27.9 ± 4.6 § | 33.5 ± 1.9 § § |
| Instilled | 235.1 ± 38.9 *** | 212.7 ± 32.3 *** | 286.8 ± 35.9 *** ‡§ |

Data are presented as mean ± standard deviation. Each group includes n = 10 animals, with 10 images per animal and 10 alveolar septum (tabique) measured per image. A two-tailed t-test was used to compare Control vs. Instilled for Observers 1 and 3. For Observer 2, a Mann–Whitney U test was applied due to the non-normal distribution of values. \*\*\*p < 0.001 compared to Control (within observer). To assess inter-observer differences, a one-way ANOVA with Tukey's post hoc test was performed for the Control group. For the Instilled group, due to non-normality in Observer 2 data, a Kruskal–Wallis test with Dunn's post hoc comparison was applied. ‡ p < 0.05 between Observer 1 and Observer 3. § p < 0.001 between Observer 2 and Observer 3.

In contrast, the 2D-ACF method, as illustrated in Figure 1, circumvents this issue by analyzing the entire image rather than relying on the measurement of an arbitrary number of isolated septum. This comprehensive approach enables global assessment of the alveolar architecture, providing a representative and unbiased metric of tissue structure. By fitting an ellipse to the 2D-ACF, we derived the *s*xy as a proxy metric, which quantifies the average spatial correlation of structural features—effectively capturing the overall septum thickness without biases (Figure 1h).

An adaptive thresholding approach based on K-means clustering was implemented to ensure consistent and unbiased identification of the ACF contour. The use of a fixed threshold—such as cutting the ACF at 1/√2 (∼0.6)—proved inadequate across diverse histological images due to variability in peak sharpness, noise, and tissue architecture. In contrast, K-means clustering allows the dynamic partitioning of ACF values into discrete intensity levels without relying on predefined assumptions (Figures 2 and 3). This approach enables effective discrimination between background noise and the central correlation peak, even in images with substantial heterogeneity, such as those in Instilled or HFD+instilled lungs. Importantly, the method adapts to image-specific features, such as the tissue-to-air ratio and full width at half maximum (FWHM) of the ACF peak, ensuring robust performance across a 400-image dataset analyzed in Figure 4. The implementation of fixed initialization parameters (e.g., random_state=42, n_init=10) guarantees reproducibility across sessions and eliminates the need for manual inspection, facilitating high-throughput and unbiased analysis.

Once the optimal contour was identified by the K-Means, an elliptical fitting procedure was applied to extract key spatial descriptors from the 2D-ACF. The selection of an elliptical model is grounded in biological relevance, as pulmonary structures often display anisotropic organization due to the directional alignment of alveolar septum, airways, and vasculature. Ellipse fitting reduces the contour complexity to five parameters—center coordinates, major and minor axes, and orientation—providing an efficient representation of spatial correlation patterns (Figure 1h). An interesting point about elliptic fitting for the 2D-ACF relies on a form of regularization, smoothing local irregularities that may arise from noise or segmentation artifacts. By focusing on the central ACF peak, this approach avoids confounding effects from disconnected contours and consistently captures the dominant spatial structure across diverse imaging conditions. The resulting σx and σy parameters offer a quantitative basis for comparing tissue anisotropy and remodeling across experimental groups.

The *s*xy parameter extracted from the 2D-ACF performed robustly across the four experimental groups (Control, HFD, Instilled, and HFD-Instilled), as shown in Figures 3 and 4. This metric not only detected significant differences between healthy and injured lungs but also sensitively captured the additive effects of HFD on lung structure (Figure 4 and Table 1). The technique’s sensitivity to subtle changes in spatial organization highlights its potential for assessing disease progression or therapeutic effects in preclinical studies. Notably, while the method assumes a level of structural homogeneity across the image, the consistent differences observed across experimental groups support its validity under controlled imaging conditions. Future work may further explore how local heterogeneity affects the autocorrelation signal and refines the method for heterogeneous tissue presentations. Another interesting aspect for future discussion is the orientation and anisotropy of the 2D-ACF. Because the lung samples were not obtained with the final aim to study spatial anisotropy, the results are difficult to interpret in an oriented tissue.

The correlation analysis between the traditional method and the automated 2D-ACF approach provides further evidence of the validity of the proposed technique (see Figure 6). In the instilled group, which exhibits substantial tissue remodeling and increased septum thickness, a strong and statistically significant correlation (r = 0.89, R² = 0.80, p < 0.001) was observed, indicating that both methods capture similar structural trends despite their methodological differences. In contrast, the control group, characterized by thinner and more homogeneous septa, showed a weaker correlation (r = 0.64, R² = 0.41, p = 0.046). These findings reinforce the strength of the 2D-ACF method, particularly under conditions of structural complexity where traditional measurements are more challenging and less reproducible. By providing consistent, unbiased quantification across experimental groups, the 2D-ACF analysis emerges as a fast, robust, and reproducible tool for studying histopathological changes in lung architecture.

While the primary focus of the present work was to evaluate the performance of the 2D-ACF analysis for quantifying alveolar septum thickness in lung tissue, the underlying principles of the method are broadly applicable to other histological contexts. Although no prior studies have applied this approach to histopathology, its successful use in analyzing the spatial distribution of mineral phases in geosciences highlights its versatility in characterizing heterogeneous structures (21). In this context, it is feasible to extend 2D-ACF analysis to hematoxylin and eosin (H&E)-stained samples from various organs. For instance, tissues exhibiting pronounced heterogeneity in nuclear size, glandular organization, or stromal density—such as in tumors, liver fibrosis, or renal pathology—could benefit from an objective spatial statistical assessment. The ability to extract geometric descriptors and quantify anisotropy or correlation length without manual segmentation makes this method particularly suitable for analyzing complex, irregular tissue architectures.

In summary, this study introduces a novel application of two-dimensional (2D) spatial autocorrelation to quantify alveolar septum thickness in histological lung images. By validating the approach across a biologically relevant experimental model of acute lung injury (ALI), which incorporates both inflammatory and metabolic stressors, we demonstrate its utility as a quantitative, observer-independent, and efficient method for assessing lung injury under challenging conditions and realistic scenarios. The 2D autocorrelation function (2D-ACF) method complements existing histopathological techniques, providing a valuable tool for future research aimed at understanding lung pathology and evaluating therapeutic interventions.

## Acknowledgments

The authors are extremely thankful to Dr. Martín Angulo for his support and valuable insights on the LPS model and manuscript discussion. ML and MJG have received an early career award (#22420230100375UD, Iniciación 2023) from the Comisión Sectorial de Investigación Científica (CSIC-Udelar). ML and MJG receive MSc and PhD fellowships from the Agencia Nacional de Investigación e Innovación (ANII) and the Comisión Académica de Posgrado (CAP-CSIC), respectively. LM is a researcher at PEDECIBA (Programa de Desarrollo de las Ciencias Básicas, Uruguay) and the Sistema Nacional de Investigadores (SNI-ANII, Uruguay). LM is supported by grants #2020–225439, #2021–240122, and #2022–252604 from the Chan Zuckerberg Initiative DAF, an advised fund of the Silicon Valley Community Foundation.

