## Supplementary Material for "Quantitative and unbiased lung alveolar septum assessment in a LPS experimental mouse model using 2D-spatial correlation image analysis from hematoxylin and eosin slides"

This supplementary material provides descriptive statistic of the data analyzed and a comprehensive overview of the methodologies, data, and additional analyses that support the findings presented in the main paper. The primary focus of this work is the development of a novel method based on the 2D ACF to estimate pulmonary edema. To further elucidate the techniques used, detailed explanations of the algorithmic approaches, including the K-Means clustering procedure employed for optimal thresholding of the ACF and additional experimental plots, figures, and programming code, are described. This will allow readers to fully understand the computational process and facilitate the replication or extension of the work.

|  | | | | | | | | |
| --- | --- | --- | --- | --- | --- | --- | --- | --- |
| **Descriptive Statistics Figure 4** | | | | | | | | |
|  | | **Ctrl** | | **Inst** | | **HFD** | | **HFD+Inst** |
| Valid |  | 10 |  | 10 |  | 10 |  | 10 |
| Mean |  | 51.8 |  | 235.4 |  | 93.9 |  | 276.8 |
| Std. Deviation |  | 9.8 |  | 24.8 |  | 29.2 |  | 48.7 |
| Coefficient of variation |  | 0.189 |  | 0.105 |  | 0.311 |  | 0.176 |
| Shapiro-Wilk |  | 0.967 |  | 0.969 |  | 0.952 |  | 0.943 |
| P-value of Shapiro-Wilk |  | 0.862 |  | 0.883 |  | 0.694 |  | 0.584 |
| Minimum |  | 35.8 |  | 200.2 |  | 52.7 |  | 175.2 |
| Maximum |  | 66.7 |  | 275.4 |  | 153.7 |  | 340.2 |
| Ctrl = Control normal diet; Inst = instilled normal diet; HFD = high fat diet; HFD+Inst = high fat diet + instilled. | | | | | | | | |

| **Descriptive Statistics Figure 5** | | | | | | | | | | | | |
| --- | --- | --- | --- | --- | --- | --- | --- | --- | --- | --- | --- | --- |
|  | | Control | | | | | | Instilled | | | | |
|  | | Obs-1 | | Obs-2 | | Obs-3 | | Obs-1 | | Obs-2 | | Obs-3 |
| Valid |  | 10 |  | 10 |  | 10 |  | 10 |  | 10 |  | 10 |
| Mean |  | 23.066 |  | 27.947 |  | 33.476 |  | 235.136 |  | 212.678 |  | 286.770 |
| Std. Deviation |  | 1.766 |  | 4.615 |  | 1.921 |  | 38.907 |  | 32.318 |  | 35.889 |
| Shapiro-Wilk |  | 0.954 |  | 0.971 |  | 0.903 |  | 0.974 |  | 0.752 |  | 0.944 |
| P-value of Shapiro-Wilk |  | 0.712 |  | 0.899 |  | 0.239 |  | 0.924 |  | 0.004 |  | 0.601 |
| Minimum |  | 20.856 |  | 20.369 |  | 30.217 |  | 179.281 |  | 185.143 |  | 209.939 |
| Maximum |  | 26.446 |  | 34.985 |  | 36.072 |  | 304.520 |  | 277.197 |  | 334.608 |

#### 1. Image preprocessing steps

**1.1 Grayscale Conversion and Gaussian Mixture Model Segmentation:**

i) Grayscale Conversion and Gaussian Blurring: The histological images are captured in RGB, but the critical information for pulmonary edema assessment lies primarily in the structural distribution of tissue and airspaces rather than color variations, so grayscale conversion simplifies the analysis while preserving this essential structural information. The application of a Gaussian blur (kernel size 11×11, σ=20) reduces noise, which is crucial for subsequent segmentation steps and smoothing of small, irrelevant details that could otherwise interfere with the autocorrelation analysis.

ii) Gaussian Mixture Model (GMM): We chose GMM with a fixed component number (k=3) over simpler thresholding methods for several critical reasons:

1. Histological images often have inhomogeneous illumination and staining, making global thresholding suboptimal
2. GMM can adaptively model multiple tissue classes (e.g., dense tissue, less dense tissue, and airspace) based on intensity distributions
3. The "tied" covariance type enforces similar shapes for the clusters while allowing different means, which better represents the physical reality of tissue variability in histological samples
4. GMM provides probabilistic assignments, which are more robust to imaging artifacts and intensity variations than hard thresholding

#### The preprocessing pipeline begins with the conversion of RGB histological images to grayscale, followed by the application of a Gaussian Mixture Model (GMM) for tissue segmentation. This approach effectively distinguishes between tissue and airspace regions.

**1.2 Small Object Removal and Hole Filling**

iii) Small White Object Removal: Small white objects (min_size=900 pixels) in histological preparations are often artifacts that do not represent the true tissue architecture and can distort the autocorrelation analysis by introducing false spatial frequencies. The chosen threshold of 900 pixels was empirically determined to remove these artifacts while preserving legitimate tissue structures effectively.

iv) Small Black Hole Filling: Small black holes are present within tissue regions (processed using binary_fill_holes), filling these holes creates more cohesive tissue regions that better represent the macroscopic tissue-airspace relationship relevant to edema assessment. This preprocessing step enhances the signal-to-noise ratio in the subsequent autocorrelation analysis by emphasizing the dominant spatial patterns while suppressing microscopic irregularities

#### To reduce noise and enhance the tissue-air interface definition, small objects were filled and small holes were removed.

```
def remove_spots(img, min_size=900):
   """Remove small white spots and fill black holes in binary image."""

   def remove_small_white_objects(binary_image, min_size=100):
       mask = (binary_image == 255).astype(bool)
       cleaned = morphology.remove_small_objects(mask, min_size=min_size)
       return (cleaned.astype(np.uint8)) * 255

   def fill_black_holes(binary_image):
       return (ndimage.binary_fill_holes(binary_image > 0)).astype(np.uint8) * 255

   max_val = img.max()
   _, thresh = cv2.threshold(img, max_val - 1, 255, cv2.THRESH_BINARY)

   cleaned = remove_small_white_objects(thresh, min_size=min_size)
   filled = fill_black_holes(cleaned)

   return filled
```

**1.3 Tissue-Air Ratio Calculation**

#### The tissue-air ratio serves as an important parameter for classifying the degree of pulmonary edema, directly quantifying a fundamental aspect of pulmonary edema - the increase in tissue volume relative to airspace. As edema increases, fluid accumulation increases the tissue component (black pixels) while reducing the available airspace (white pixels). By calculating this ratio first, it can adaptively select the most appropriate ACF threshold level for each image, enhancing both sensitivity and specificity of the method. This metric is robust and straightforward to calculate from the binary segmentation, providing a valuable first-order parameter without complex computation.

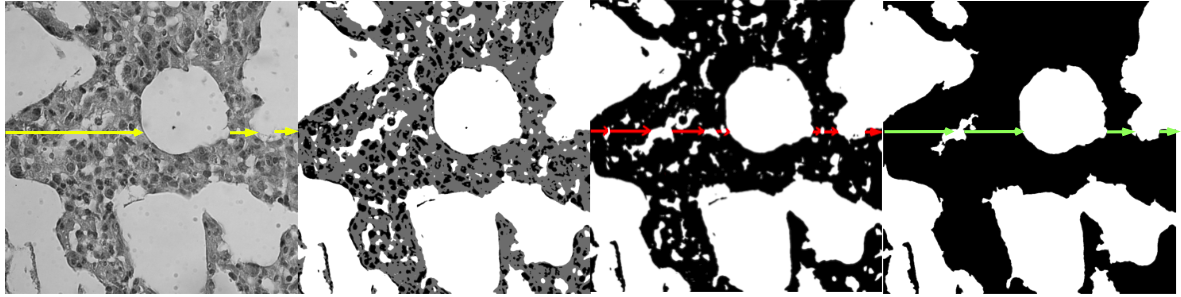

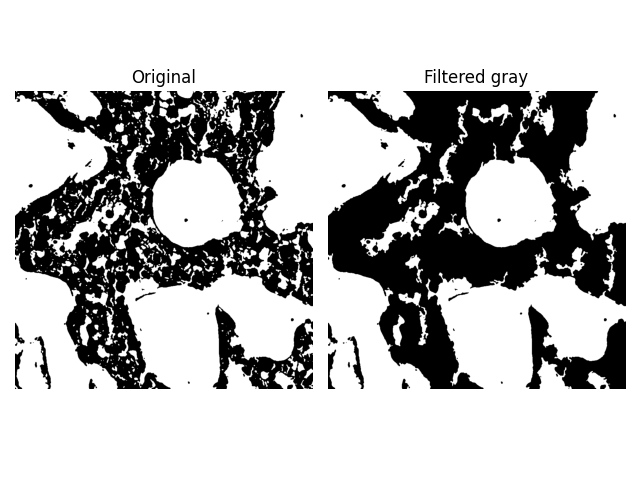

Figure image of the instilled group without applying the GMM technique, only using OTSU's Threshold to automatically find the optimal threshold value, as can be seen, the result is an image with a lot of blank spaces or gaps.

#### 2. 2D Autocorrelation Function Calculation

The Fourier transform has been employed as a method to calculate the autocorrelation function rather than direct spatial domain computation for the next reasons:

- Computational Efficiency: For large images (1024×1024), the FFT-based approach has a complexity of O(N²log(N²)) compared to O(N⁴) for direct spatial domain calculation.
- Numerical Stability: The FFT approach avoids cumulative rounding errors that can occur in direct spatial calculations.
- The power spectral density (|FFT|²) captures the energy distribution across spatial frequencies. This transformation emphasizes dominant structural patterns while de-emphasizing random noise, making it ideal for detecting the characteristic spatial scales of tissue architecture.

The min-max normalization (ACF-min)/(max-min) ensures that all ACF values fall within [0,1], facilitating consistent thresholding across images and different images can be directly compared despite variations in absolute intensity values

Finally the fftshift operation centers the zero-frequency component, which aligns the ACF with natural spatial coordinates for more intuitive interpretation, facilitates measurement of correlation distances from the central maximum and simplifies subsequent contour detection and fitting operation.

```
def calcule_2D_ACF(img, apply_window):
   if apply_window:
       hann = np.hanning(n)[:, None] * np.hanning(n)
       img = img * hann
       fft_img = np.fft.fft2(img, s=(2048, 2048))
       psd = np.abs(fft_img) ** 2
       acf = np.fft.ifft2(psd).real
       acf = np.fft.fftshift(acf)
       acf = acf[n - n // 2:n + n // 2, n - n // 2:n + n // 2]
       ACF = (acf - acf.min()) / (acf.max() - acf.min())
   else:
       fftn = np.fft.fft2(img)
       psd = np.abs(fftn) ** 2
       ACF = np.fft.ifft2(psd).real
       ACF = (ACF - ACF.min()) / (ACF.max() - ACF.min())
       ACF = np.fft.fftshift(ACF)
   labels, best_levels = apply_kmeans_clustering(ACF, n_clusters=5)
   return ACF, best_levels
```

#### 3.Threshold Selection Logic

**3.1 Justification for using K-Means clustering to identify optimal ACF threshold levels**

The choice of K-Means clustering for determining the appropriate levels at which to cut the 2D autocorrelation function is motivated by the need for an adaptive, data-driven approach that accounts for variations in the structure of different histological images. The previous method used where cut the peak at a fixed level (e.g: 1/sqrt(2)≈ 0.6), failed to generalize across different images due to variations in ACF peak sharpness, noise levels, and tissue distribution patterns. The ACF exhibits distinct intensity regions that correspond to different structural patterns in the images. By applying K-Means clustering, one can automatically group ACF values into K discrete levels without relying on predefined thresholds. This approach allows to distinguish between background noise and meaningful structure regions. Identify the most relevant contour level where the central ACF peak retains an elliptical shape without requiring prior knowledge of tissue-specific thresholds, making it robust across different imaging conditions and pathological presentations.

Adaptively adjust the threshold based on image characteristics rather than using a fixed approach.
Through empirical observations, it has been found that the Tissue/Air Ratio (black/white pixel ratio in the original image) and the full-width at half maximum (FWHM) of the ACF peak influence the optimal level selection. K-Means clustering provides an efficient way to categorize the ACF values, making it possible to select an appropriate threshold dynamically, ensuring that threshold selection is consistent and reproducible across a dataset of 400 images. In addition K-Means is computationally efficient and can be applied in batch mode to large datasets by using this clustering technique, eliminate the need for manual inspection, facilitating high-throughput image analysis. The fixed random state (random_state=42) and multiple initializations (n_init=10) ensure consistent cluster identification across different analysis sessions.

Implementation Details:

The clustering algorithm processes flattened ACF values to identify four distinct clusters (n_clusters=4), corresponding to the natural progression from normal aerated tissue to severe consolidation. The sorted cluster centers provide ordered threshold values that align with increasing tissue density. The 4-cluster configuration was selected based on the clinical classification of pulmonary edema severity and validated against radiological assessments.

```
def apply_kmeans_clustering(acf: np.ndarray, n_clusters: int = 4) -> Tuple[np.ndarray, np.ndarray]:
   """Apply KMeans to cluster ACF values."""
   values = acf.flatten().reshape(-1, 1)
   kmeans = KMeans(n_clusters=n_clusters, random_state=42, n_init=10)
   kmeans.fit(values)
   labels = kmeans.labels_.reshape(acf.shape)
   centers = np.sort(kmeans.cluster_centers_.flatten())
   return labels, np.round(centers, 2)
```

**3.2 Limitations of Fixed Thresholding at 0.6:**

Initially, a fixed ACF threshold of 0.6 was applied to determine the contour for further analysis. However, this approach has several drawbacks:

- Lack of Adaptability: The ACF intensity varies significantly across different lung images due to differences in tissue structure and imaging conditions. A single fixed threshold does not account for these variations.
- Loss of Elliptical Shape: In some cases, applying a strict 0.6 threshold results in distorted contours that do not resemble an ellipse, leading to inaccurate measurements.
- Inconsistent Interpretation: For images with sharper ACF peaks, the fixed threshold cuts too high, reducing the effective region for analysis. For images with broader peaks, the fixed threshold cuts too low, including noise.

Now instead of assuming a universal 0.6 cut-off, K-Means groups ACF intensity values into K clusters, allowing to select the most meaningful level based on the image characteristics. Cluster 0 is assumed across all the dataset as a noise signal.

### 3.3 Threshold Selection

The selection of the appropriate threshold for contour extraction is a critical step in our method. A decision algorithm was implemented that considers both the tissue-air ratio and FWHM values. The complexity of pulmonary edema manifestation necessitates a multi-parameter approach.

The FWHM of the ACF peak provides a direct measurement of the characteristic length scale of tissue structures. This metric is physically meaningful because in normal lung tissue, the sharp transition between tissue and airspace creates rapid correlation decay and thus a narrow ACF peak (small FWHM) and in edematous tissue, the more gradual tissue density variations lead to extended spatial correlation and a broader ACF peak (large FWHM).

Different stages of edema require different analytical approaches. The algorithm dynamically selects the optimal cluster based on both tissue-air ratio and Full Width Half Maximum (FWHM) values.

Selection Criteria:

- Severe Edema (tissue-air ratio ≥ 5.0): select cluster 1 to capture fine structure within predominantly solid tissue areas.
- High tissue-air ratio with moderate FWHM (tissue-air ratio > 4.0): select cluster 1 for severe consolidation cases.
- Moderate-to-High Edema (tissue-air ratio > 2.5): select cluster 1, 2, or 3 based on specific tissue-air ratio thresholds and FWHM values, with cluster 2 preferred for ratios 3.0-4.0 and cluster 3 for borderline cases with small FWHM.
- Mild-to-Moderate Edema (1.0 ≤ tissue-air ratio ≤ 2.5): select cluster 2 or 3 based on tissue-air ratio and FWHM characteristics, with cluster 2 for ratios ≥ 2.0 and cluster 3 for ratios 1.0-2.0.
- Normal-to-Minimal Edema (tissue-air ratio < 1.0): select cluster 3 or 4 based on FWHM, with cluster 4 preferred for very low ratios (≤ 0.6) and cases with large FWHM values, optimizing for sharp transitions between tissue septum and airspaces.

The adaptive selection algorithm implements a hierarchical decision tree that considers both tissue-air ratio and FWHM values:

Extreme conditions: Direct cluster assignment for very low (≤ 0.6) and very high (≥ 5.0) tissue-air ratios

FWHM-based stratification: Different thresholds applied based on FWHM relative to image dimensions (n/4, n/8, n/16, n/10)

Transition zones: Gradual cluster selection for intermediate tissue-air ratios (1.0-3.0 range) with FWHM modulation

Edge case handling: Specific logic for borderline cases where tissue-air ratio and FWHM suggest different optimal clusters.

```
def select_best_cluster(fwhm_value, tissue_air_ratio):

   if tissue_air_ratio <= 0.6: return 4
   if tissue_air_ratio >= 5.0: return 1
   if fwhm_value > n // 4:
       return 4 if tissue_air_ratio <= 1.0 else 3 if tissue_air_ratio <= 2.0 else 2
   if fwhm_value >= n // 8:
       return 4 if tissue_air_ratio < 0.9 else 3 if tissue_air_ratio <= 2.0 else 2
   if tissue_air_ratio > 2.5:
       return 1 if tissue_air_ratio > 4.0 else 2 if tissue_air_ratio > 3.0 or fwhm_value < n // 16 else 3
   return 2 if tissue_air_ratio >= 2.0 else 3 if tissue_air_ratio >= 1.0 or fwhm_value > n // 8 else 4 if n // 10 > fwhm_value else 3
```

#### 4. Elliptical Contour Fitting

It has been chosen an elliptical model for contour fitting for several compelling reasons:

1. Biological Relevance: Pulmonary structures often exhibit directional preferences due to the anatomical organization of airways and blood vessels, which is well-captured by an elliptical model.
2. Parameter Reduction: The ellipse reduces the complex contour to just five parameters (center x, center y, major axis, minor axis, orientation), enabling efficient quantitative comparison between samples.
3. Robustness to Noise: The elliptical fit essentially performs a form of regularization, smoothing out irregular contour variations that might result from noise or artifacts.

Specifically target the contour that surrounds the ACF center because this contour corresponds to the primary correlation structure of interest, ensures consistency across images with different characteristics and it avoids the complexity of dealing with multiple disconnected contours that may appear at certain threshold levels

Contour Validation: is_valid_curved_shape function implements multiple validation criteria:

1. Aspect Ratio Constraints: Extreme aspect ratios (outside 1.0-2.0) often indicate problematic fits or anisotropic artifacts
2. Solidity Threshold: The solidity (ratio of contour area to convex hull area) measures the "smoothness" of the contour. Values below 0.65 suggest irregular shapes that may not be well-represented by an ellipse.
3. Fit Error Tolerance: By comparing the actual contour area to the theoretical ellipse area, we ensure that the elliptical model adequately represents the underlying data. The tolerance of 0.30 (30%) was empirically determined to balance fidelity with robustness.

Adaptive Threshold Adjustment: If the initial contour fails the validation criteria, then incrementally adjust the threshold (by delta=0.02) until a valid contour is found. This adaptive approach ensures that even challenging images yield meaningful measurements

```
def fit_contours(g: np.ndarray, n: int, file_name: str, thresh_level: float) -> Tuple[float, float, float]:
   """Fit an ellipse to the central contour of a thresholded ACF image."""
   center = (n // 2, n // 2)
   delta = 0.001 if thresh_level < 0.1 else 0.01

   while True:
       _, binary = normalize_and_threshold(g, thresh_level)
       cnt = get_central_contour(binary, center)
       if len(cnt) < 5:
           thresh_level += delta
           continue

       ellipse = cv2.fitEllipse(cnt)
       _, (w, h), _ = ellipse

       if is_valid_ellipse(cnt, n) and not (0.5 * h >= n / np.sqrt(2)) and h < n:
           break

       thresh_level += delta

   return round(w, 4), round(h, 4), ellipse[2]
```
